# High-resolution *Staphylococcus* profiling reveals intra-species diversity in a single skin niche

**DOI:** 10.1101/2025.04.01.646391

**Authors:** Reyme Herman, Daniel T. West, Sean Meaden, Imran Khan, Robert Cornmell, Joanne Hunt, Michelle Rudden, Holly N. Wilkinson, Matthew J. Hardman, Anthony J. Wilkinson, Barry Murphy, Gavin H. Thomas

## Abstract

The skin microbiome is dominated by a few key genera, among which *Staphylococcus* is one of the most well-characterised. Recent studies have examined the roles of various *Staphylococcus* species such as *S. epidermidis* and *S. hominis* within broader skin microbial communities. However, these investigations often rely on isolates from multiple individuals and hence, limiting their ability to capture intra-community interactions. In this study, we focused on the axillary microbiome of a single healthy individual to characterise the genetic and functional diversity of resident *Staphylococcus* isolates. Using a low-cost, high-throughput DNA extraction and long-read whole genome sequencing pipeline, we generated complete genomes for 93 isolates spanning seven genetically distinct lineages across three major skin species. These comprised of one dominant with three additional lineages of *S. epidermidis*, two of *S. hominis* and one of *S. capitis*. Functional and metabolic analyses revealed species- and strain-specific features, suggesting potential metabolic cross-feeding and specialisation within this community, including within strains of *S. epidermidis*. These findings highlight the metabolic complexity and potential interdependence of staphylococci inhabiting a single skin site and the need for strain-level resolution of the community. The strains form part of the York Skin Microbiome (YSM) collection, a growing open biobank of genetically diverse skin isolates from matched individuals.

## Introduction

Human skin is home to a wide range of microorganisms that have evolved with their host over many millions of years (1–3). Naturally, these organisms interact with each other and the host in a symbiotic relationship to establish a favourable environment which protects the host while allowing for these microorganisms to thrive in generally nutrient limiting conditions. The multiple skin sites of the human body have some characteristic differences, such as being oily/sebaceous (e.g. face), dry/sebaceous poor (e.g. volar forearm) or moist (e.g. underarm/axilla) which then dictates the abundance and the diversity of microbes in the microbiome. The genus *Staphylococcus* is one of the most abundant on human skin and can be found in multiple skin environments. While some species within this genus are well-known human pathogens (e.g. *Staphylococcus aureus*), cutaneous species of *Staphylococcus* play multiple roles in maintaining skin health. For example, the commonly identified species *S. epidermidis* expresses a sphingomyelinase which contributes to the production of ceramides which is important in maintaining the epithelial barrier (4). Another study identified strains of *S. epidermidis* that produce 6-N-hydroxyaminopurine which reduces the occurrence of ultraviolet-radiation induced cutaneous tumours in mice (5). Recent studies have also demonstrated the various roles for staphylococci have in modulating the immune response. For example, *S. epidermidis* is able to interact with dendritic cells to activate IL-17A^+^ CD8^+^ T-cells to aid in the protection against skin pathogens (6,7). Rather than priming the host immune system, multiple cutaneous staphylococci are also able to directly suppress known skin pathogens by producing anti-microbial peptides (e.g. *S. hominis*: hominicin (8), *S. capitis*: capidermicin (9) and *S. lugdunensis*: lugdunin (10)). While the role of staphylococci in maintaining skin health is evident, species within this genus can also participate in ancient processes like malodour production. We previously identified a monophyletic group of coagulase-negative staphylococci (CoNS) including *S. hominis* and *S. lugdunensis* which are able to metabolise the human apocrine gland derived metabolite, Cys-Gly-3M3SH, and convert it to an odorous molecule we associate to human malodour (3,11–13). Interestingly, different isolates of the same species of *Staphylococcus* can have varying levels of activity against Cys-Gly-3M3SH suggesting as yet undiscovered genetic features that could control malodour formation (11,14). In this study, we isolated the axillary staphylococci sampled from a single individual to understand the different genetic components that form the *Staphylococcus* community in a distinct body site. We adapted a low-cost high throughput DNA extraction and whole genome sequencing pipeline to enable the genetic assessment of these isolates. Through this, we reveal the genetic differences among these *Staphylococcus* isolates which could contribute to maintaining the skin microbiome community and ultimately skin health. This collection of staphylococci contributes to the establishment of the <u>Y</u>ork <u>S</u>kin <u>M</u>icrobiome (YSM) isolate resource of genetically diverse isolates from a variety of genera from matched individuals.

## Results

### Isolation and genetic characterisation of axillary Staphylococcus

The axilla of the volunteer was swabbed and then the captured material was plated onto a selective agar to enrich for cutaneous staphylococci. 96 isolates were harvested and stocked in the York Skin Microbiome (YSM) collection. The genomic DNA from these isolates was then extracted for long-read whole genome sequencing using long-read sequencing on the PromethION platform. We generated complete genome sequences together with accompanying plasmid sequences for 93 of the 96 isolates. Using Multilocus Sequence Typing (MLST) (15), 90 of these isolates were determined to be *S. epidermidis* and 3 were *S. hominis* (Supplementary Tables 1 and 2). To determine the diversity of these isolates, we generated a Hadamard matrix (Fig 1A) using the pairwise average nucleotide identities and coverage of each alignment calculated on pyani (16). The 90 *S. epidermidis* isolates are grouped into 4 distinct clusters (SE1, SE2, SE3 and SE4) with isolates of SE1 dominating this population of *S. epidermidis* isolates. The 3 *S. hominis* isolates were found to be from two separate clusters (SH1 and SH2). In a parallel experiment involving corynebacteria from the same individual (17), we identified 27 additional isolates of staphylococci using corynebacteria enriching media containing Fosfomycin. Analysis using pyani suggested these isolates are genetically highly similar and hence are grouped in the same cluster (S) (Fig 1A). Unsurprisingly, data from Fig 1A suggest we have redundancies in the library of *Staphylococcus* isolates from the volunteer due to indiscriminate large-scale isolation. Hence, we performed dereplication of the isolates using drep (18) with a secondary cluster threshold of 99.5% to define a set of 7 genomes to represent isolates from clusters SE1, SE2, SE3, SE4, SH1, SH2 and S for use in downstream analysis. The core genome of the 7 dereplicated isolates and type strains of other skin associated staphylococci were derived and aligned using Roary (19). The resulting alignment was then used to derive a maximum-likelihood tree (Fig 1B). The core genome of isolate YSMAA1_1_H3st, representing isolate cluster S (Fig 1A), clustered with *S. capitis* and hence was assigned the species. The full 16S sequences of YSMAA1_1_H3st and *S. capitis* DSM 6717 were found to be identical, further confirming the species assignment (Supplementary Fig 1). This approach provided us with complete genomes of multiple genetically different staphylococci (4 *S. epidermidis*, 2 *S. hominis* and 1 *S. capitis*) from the same individual which enables us to understand how these bacteria may have interacted in the microbiome they were isolated from.

**Fig 1.**
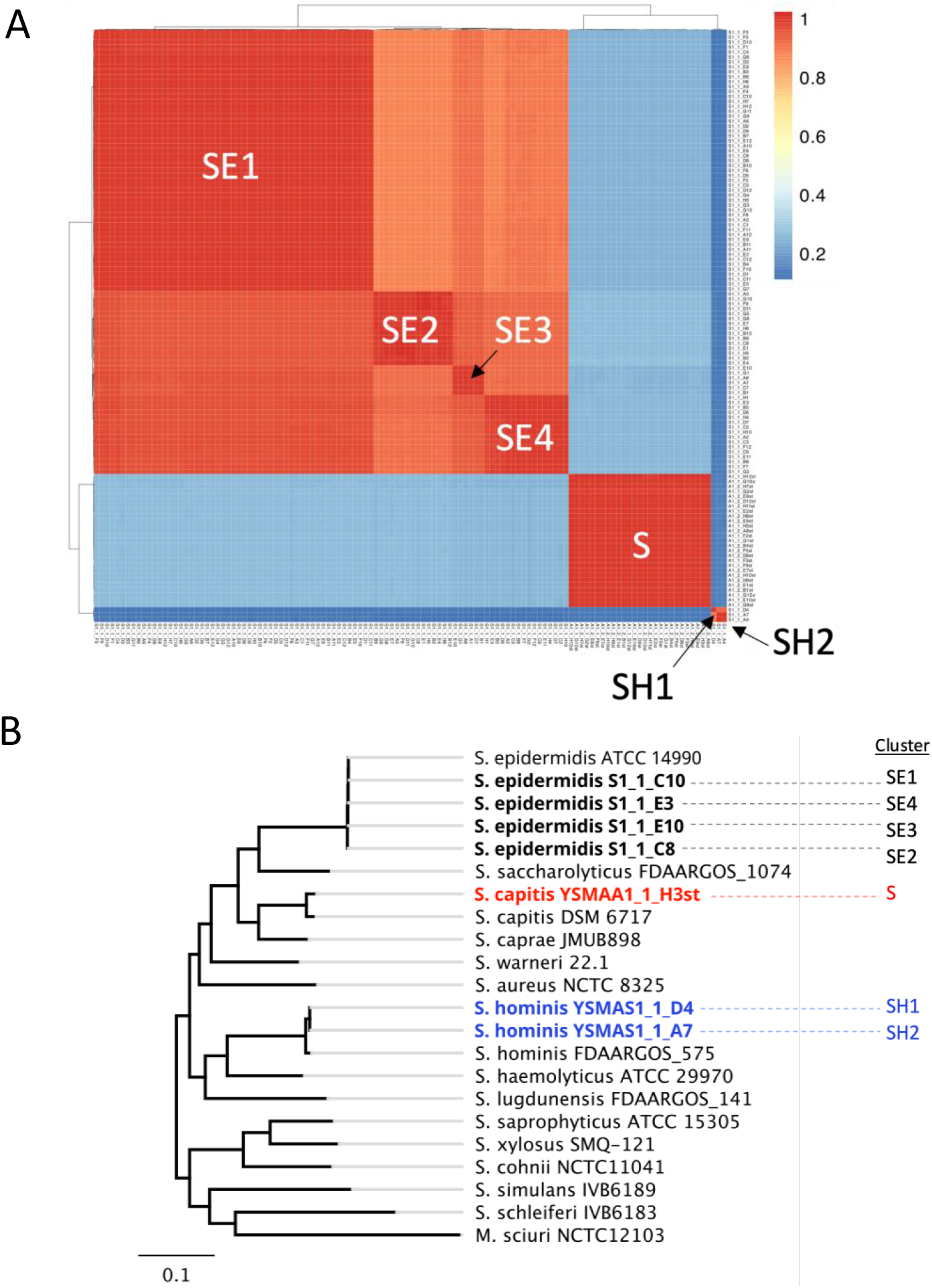
Diversity of 120 axillary staphylococci isolates. (**A**) A Hadamard matrix was constructed using the product of the pairwise-average nucleotide identities and the pairwise-alignment coverage and plotted using the pheatmap package on R. Isolate clusters were identified as SE1-4, SH1-2 or S. (**B**) Core genome maximum likelihood tree of all 7 dereplicated isolates with type strains of known cutaneous staphylococci. Clusters from (A) were identified next to the respective isolates and colour coded by species.

### Species- and isolate-specific genetic traits reveal potential roles and functional networks within a single environment

Different human body sites like the skin are thought to be an environment with networks of interactions between resident organisms of the microbiome and the host. Multiple synergistic and antagonistic relationships exist in these niches which helps maintain a healthy environment for the host. We performed a pangenome analysis using anvi’o (20) (Fig 2A) which allowed us to identify genes specific to each species as well as genes only found in individual isolates (singletons) with the aim of revealing the roles of each genetically distinct isolate in this environment.

**Fig 2.**
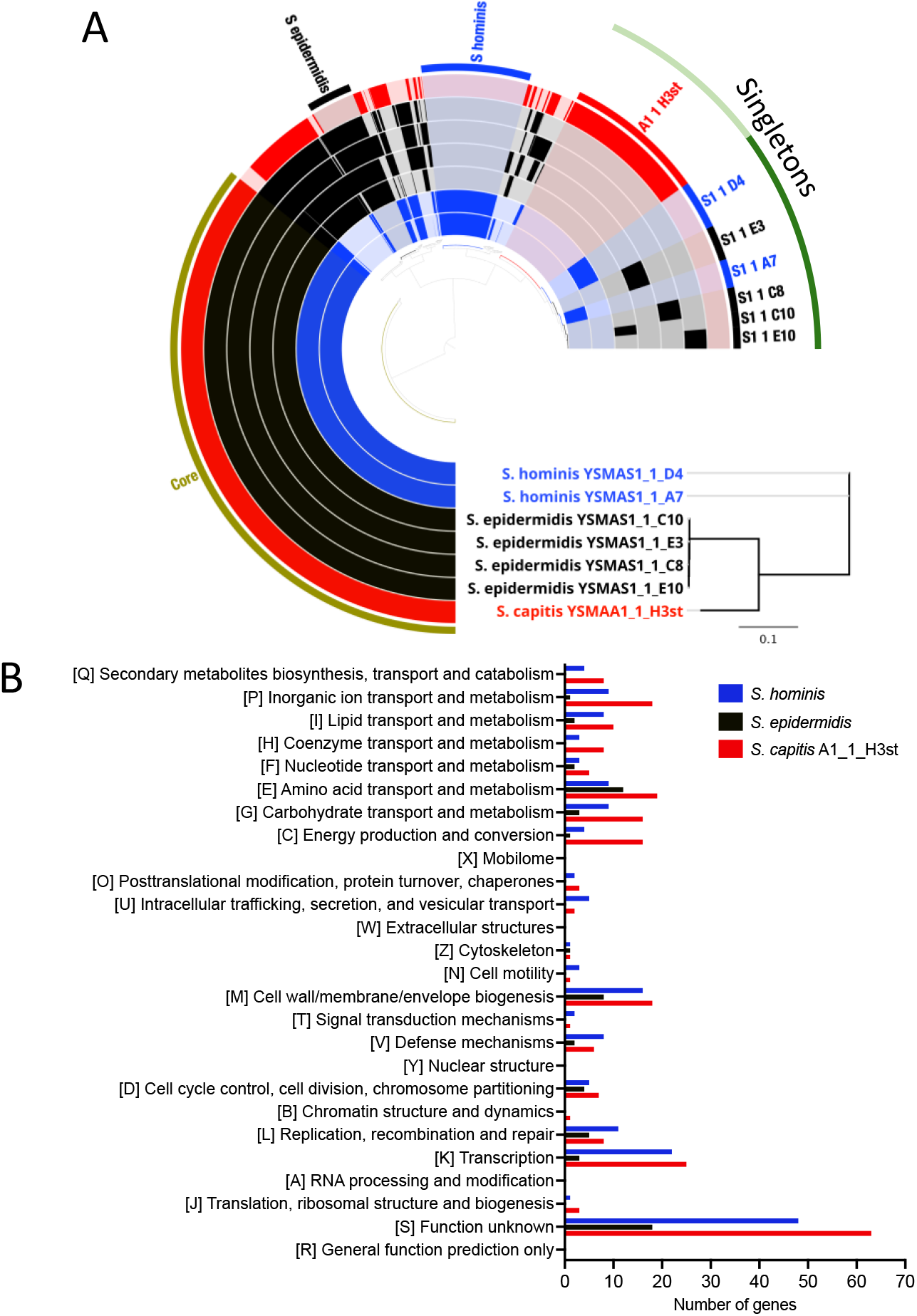
(**A**) Pangenome analysis of the 7 dereplicated axillary staphylococci isolates identified species-specific clusters (*S. epidermidis* and *S. hominis*) and gene clusters belonging to only one isolate (singletons). The A1 1 H3st gene cluster identified in *S. capitis* YSMAA1_1_H3st was used in species-specific comparisons. The pangenome was calculated and visualised on anvio-8. (**B**) COG categories were assigned to species-specific gene clusters using eggNOG-mapper and visualised on Graphpad Prism 10.

The pangenome analysis identified 1609 core genes shared by all tested isolates of this genus with an additional 116 core *S. epidermidis* genes and 281 core *S. hominis* genes. These genes were assigned putative functions (COG) using eggNOG-mapper (21,22) to help elucidate the roles of the various species in the underarm of this single individual. As we were only able to isolate one subtype of *S. capitis*, we included the singletons identified for isolate *S. capitis* YSMAA1_1_H3st to represent the species. Major species-level differences were in the number of genes predicted to be involved in nucleotide related processes like transcription (K) and replication, recombination and repair (L) and also cell surface biogenesis (M) (Fig 2B). We also observed significant differences in the number of genes predicted to be involved in transport processes and metabolism, suggesting there may be species specific pathways. To further resolve the diversity of these isolates, we performed the same functional analysis on the singletons of all *S. hominis* and *S. epidermidis* isolates and compared them to other isolates of the same species (Fig 3). Even within species level comparisons, we observe the largest differences in transcription (K) and replication, recombination and repair (L). We observed fewer differences between isolates of the same species for metabolism related genes.

**Fig 3.**
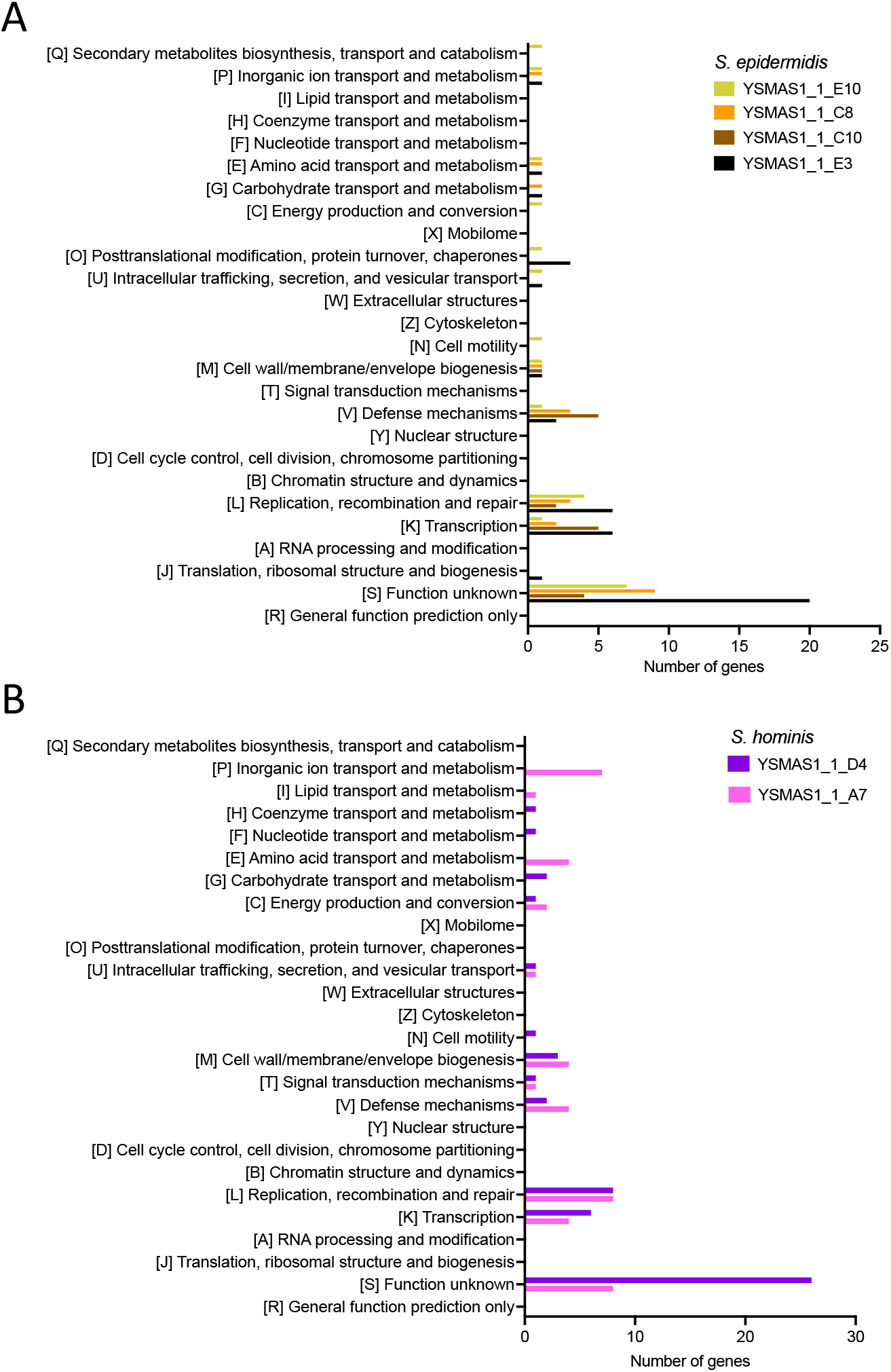
COG categories were assigned to singletons from (**A**) *S. epidermidis* and (**B**) *S. hominis* isolates using eggNOG-mapper and visualised on Graphpad Prism 10.

We also performed KEGG analysis on all 7 isolates to map full pathways to complement the COG analysis, in particular to assess the presence of known metabolic pathways (Fig 4). As observed in the COG analysis, there seem to be some correlation between the presence of some complete pathways and specific species. Here, we discuss some examples of the species- and subspecies/strain-level differences which could allude to potential metabolic functions in the microbiome. Within the amino acid metabolism pathways, the ornithine biosynthesis module (M00028) seems to only be complete in *S. epidermidis* isolates and the entire module was found to be missing from *S. hominis* and *S. capitis*. Ornithine can be used in the biosynthesis of arginine as described in the KEGG module M00844 which is found to be present in all 7 isolates (Fig 4). For this purpose, ornithine could be imported from the environment using the ornithine transporter ArcD (23). While the gene encoding for ArcD was identified in *S. capitis* YSMAA1_1_H3st, this transporter was not identified in the genomes of either of the two *S. hominis* isolates, suggesting the presence of a different ornithine transport pathway.

**Fig 4.**
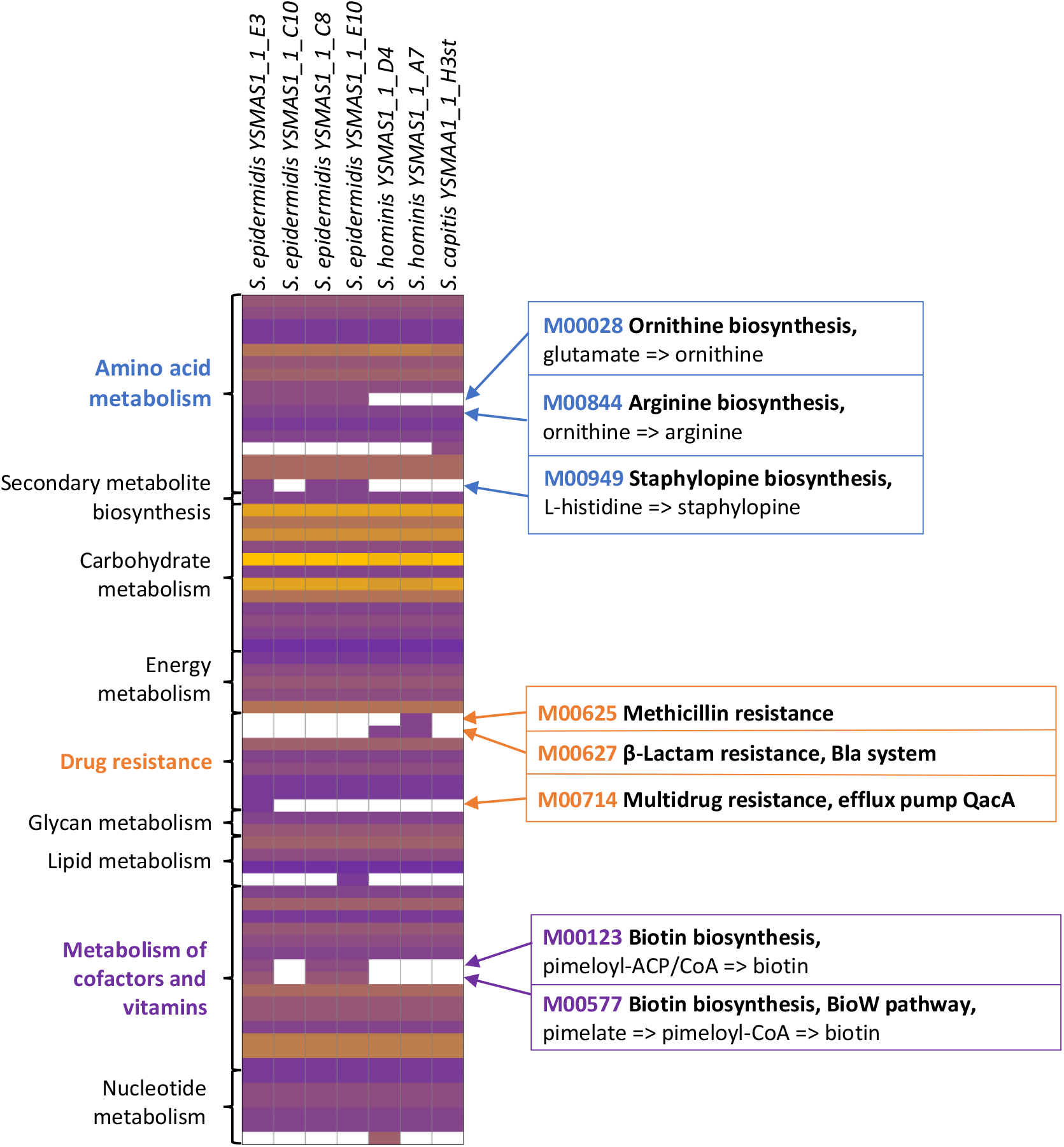
Metabolic pathways in genetically distinct axillary staphylococci reveals species and isolate level variation within the isolates. Each complete KEGG module for each isolate were placed on a heatmap where the modules with the largest number of reactions are in yellow and lowest in purple with absent modules in white. Modules of interest were described on the right of the heatmap. KEGG modules were grouped according to the categories on the left of the heatmap.

From this analysis, we also observed differences within isolates of individual species. For example, the staphylopine biosynthesis pathway (M00949), deriving the zinc-binding siderophore from histidine (24), is found in three *S. epidermidis* isolates. *S. epidermidis* YSMAS1_1_C10, both *S. hominis* and the *S. capitis* isolates were missing all the genes required for staphylopine biosynthesis. Rather than synthesising staphylopine, these isolates could instead utilise and import this siderophore, however, they lack the direct orthologue of the *S. aureus* staphylopine CntABCDF ATP-binding cassette (ABC) permease (25).

Similarly, the biotin biosynthesis pathways (M00123 and M00577) seem to only be present in *S. epidermidis* but not in *S. epidermidis* YSMAS1_1_C10. These biotin biosynthetic pathways are absent in *S. hominis* and *S. capitis* isolates which suggests some subtypes of cutaneous *S. epidermidi*s could be the source of biotin for other auxotrophic species of this genus in the same community. The transporter BioY has been identified as a probable biotin uptake system in *S. aureus* (26) with the gene encoding BioY identified in all 4 isolates deficient in biotin biosynthetic pathways. Using the complete genomes of these isolates, we were able to perform an in-depth metabolic analysis to reveal the potential metabolic cross-talk between members of the same skin microbiome.

In addition to metabolism, we also assessed the genomes for resistance cassettes and defence systems to understand the inherent protective measures which could maintain the community of staphylococci from the same site. In the KEGG analysis (Fig 4) we observed the presence of modules providing resistance to antimicrobials including β-lactams via the bla system (M00627) and the mec systems (M00625) (27,28) in specific *S. hominis* isolates and to some antiseptics via the efflux pump QacA (29) in *S. epidermidis* YSMAS1_1_E3. These findings were confirmed using ResFinder (Supplementary Table 3).

The COG analysis (Fig 3) also identified species- and isolate-specific anti-phage defence systems (X) which suggest that we can expect an array of different systems in these isolates. We then analysed the genomes of these isolates on padloc (30) to reveal putative defence systems (Fig 5). As expected, the padloc analysis predicted the presence of a wide range of defence systems. Some systems are found in all isolates of the same species like the Type I and IV restriction modification (RM) systems in S. *epidermidis* isolates and the Type II RM system in *S. hominis* isolates.

**Fig 5.**
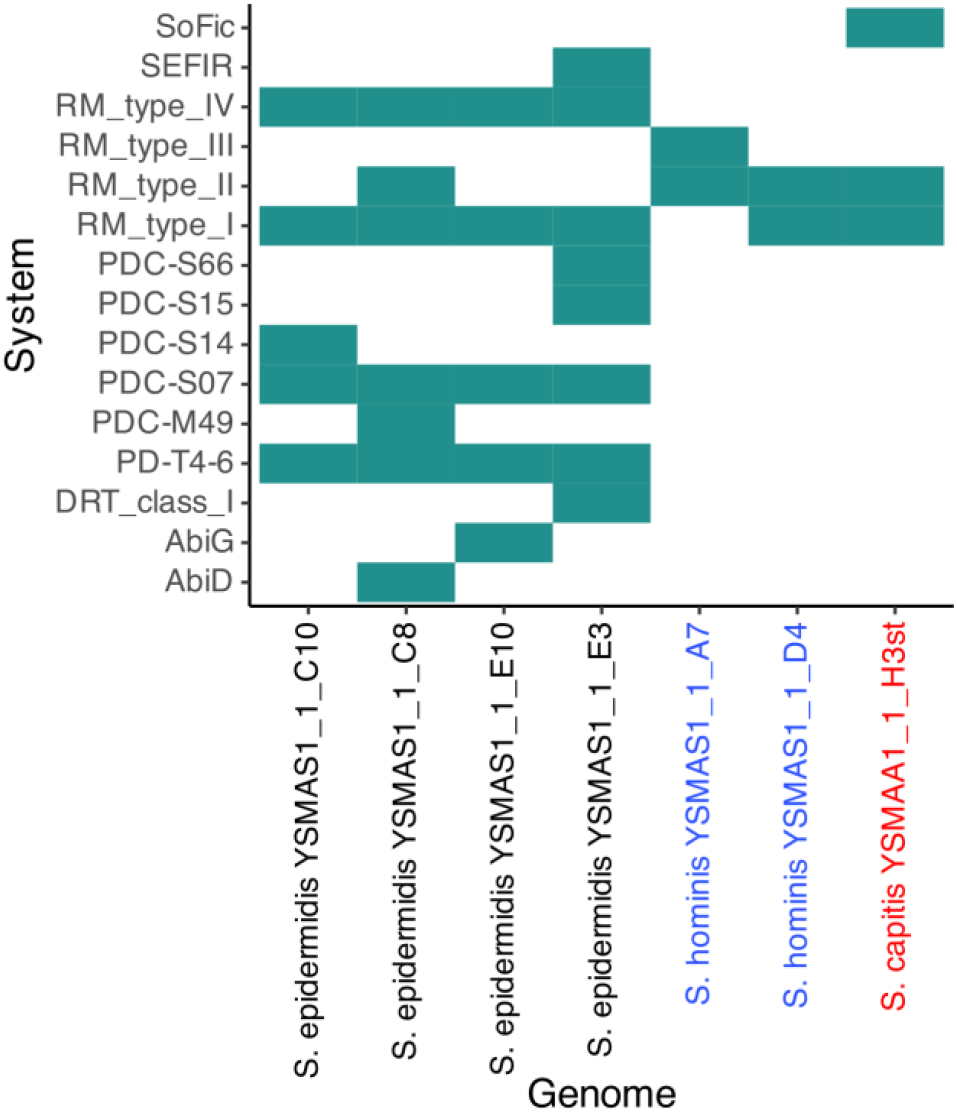
Phage defence systems were predicted in the 7 dereplicated genomes using padloc. Only one of each system was identified in each relevant genome. Names of isolates were coloured according to the species they belong to (*S. epidermidis*: Black, *S. hominis*: Blue, *S. capitis*: Red). Data was visualised using R.

However, as suggested by the species-specific COG analyses (Fig 3), we also see isolate specific systems. Notably, in *S. epidermidis*, we observed the presence of the Type II RM system, the Type I defence-associated reverse transcriptase (DRT_class_I), the abortive infection related systems (AbiG and AbiD) and SEFIR in either of the 4 dereplicated isolates. Further, we also identified putative defence systems including some previously reported by Vassallo *et al*. (31) but these were not found in both *S. hominis* isolates and *S. capitis* YSMAA1_1_H3st. In *S. hominis*, predicted defence systems were limited to RM systems with the isolates either possessing a Type III or a Type I RM system alongside the Type II RM system. We could use information from this analysis to aid in the design of genetic tools to target specific species or even specific isolates which will be invaluable in the understanding the communities of the skin microbiome.

## Discussion

Recent advances in sequencing technologies have played a crucial role in revolutionising our understanding of the human skin microbiome. Both short- and long-read sequencing technologies have provided us with the means to unlock the microbiome to study the abundance of different species in a single community through metagenomics and also enable cheaper whole genome sequencing of isolates. In this study, we have proposed the use of a staphylococci-focused isolation approach involving the use of a low-cost total DNA extraction procedure and whole genome sequencing for the isolation and characterisation of cutaneous species. We tested this approach using isolates derived from a single volunteer which allowed us to assemble a library of 120 staphylococcal isolates containing 7 genetically distinct lineages of isolates from three different species.

Through genomic analysis and downstream functional annotation, we identified potential roles for isolates of three major cutaneous *Staphylococcus* species in maintaining the skin microbiome and promoting overall skin health. The subspecies-level resolution of the KEGG analysis allowed us to identify potential metabolic networks involving molecules like ornithine, staphylopine and biotin, indicating possible cross-talk across different species (Fig 4). However, our results may be influenced by limitations in current isolation methods. Our protocols yielded a disproportionate number of *S. epidermidis* isolates, potentially underrepresenting other *Staphylococcus* species. To address this, future studies could explore the design of targeted enrichment media aimed at improving species diversity. Insights from the KEGG analysis (Fig. 4) could inform the development of such media, demonstrating the practical utility of this genomic dataset beyond descriptive analysis.

Our workflow, powered by Oxford Nanopore’s long-read whole genome sequencing technology, provided us with the ability to genetically characterise the library of isolates at a low per sample cost and could potentially be adapted to other laboratory setups. The methods and findings of this study provide a platform to establish larger scale studies of volunteers and isolates of their skin microbiome, not just on healthy individuals but could also be adapted for studies on diseased skin. The strain-level resolution described by our approach provides unparalleled access to fine-scale genetic variation, offering a powerful tool for dissecting microbe–microbe and microbe–host interactions. Ultimately, such insights will be critical for understanding how specific microbial strains contribute to skin health and disease, and for identifying potential targets for therapeutic modulation of the skin microbiome.

## Materials and methods

### Sampling procedures

The adult male volunteer was recruited for axillary swabbing at Unilever R&D, Bebington, UK. The volunteer was in good general health, had not used any cosmetic products for 24 hours before the study, was not taking antibiotics, had no active skin conditions, has not suffered from eczema in the last 5 years nor have they ever had psoriasis. Axillary swabs were collected by swabbing in a linear motion 20 times using eSwabs (Copan) on both underarms and then placed into Amies transport medium.

### Bacterial culturing

Swabs were plated onto SS solid media (10 g L^−1^ tryptone, 5 g L^−1^ Lemco powder, 3 g L^−1^ yeast extract, 13 g L^−1^ agar no. 1, 10 g L^−1^ sodium pyruvate, 0.5 g L^−1^ glycine, 22.5 g L^−1^ potassium thiocyanate, 1.2 g L^−1^ disodium hydrogen orthophosphate dihydrate, 0.67 g L^−1^ sodium dihydrogen phosphate, 2 g L^−1^ lithium chloride, 10 mL L^−1^ glycerol, pH 7.2) or ACP solid media (39.5 g L^−1^ Blood Agar base no. 2, 3 g L^−1^ yeast extract, 2 g L^−1^ glucose, 5 mL L^−1^ Tween 80, 50 mL L^−1^ defibrinated horse blood, 100 mg L^−1^ fosfomycin) to enrich for staphylococci and corynebacteria respectively. Colonies were allowed to grow for up to 2 days at 37°C. Colonies were picked and grown overnight in brain-heart infusion broth + 1 % Tween-80 (BHIT) at 37°C, shaking at 200 rpm. Cultures were stocked in BHIT supplemented with 10 % glycerol in 2 mL 96 well plates.

### DNA extraction

Total (genomic and plasmid) DNA extraction was performed using a previously reported procedure (17) with the following changes: the enzymatic lysis buffer used also included 125 μg/mL lysostaphin, the enzymatic lysis step was performed over 5 hours and DNA precipitation was performed for 15 minutes at room temperature.

### DNA library preparation, whole genome sequencing and assembly

The modified Oxford Nanopore Native Barcoding Kit 96 V14 DNA library preparation protocol reported in Herman *et al*. (17) was used. The prepared library was loaded into a PromethION R10.4.1 flow cell and sequencing was performed for 72 hours and base-called using the singleplex high-accuracy model, 400 bps on MinKNOW 23.07.5. The reads were assembled with the EPI2ME Labs wf-bacterial-genomes isolate workflow (Flye (32), Medaka (Oxford Nanopore Technologies), Prokka (33).

### Bioinformatics tools

Pairwise genome comparisons of all assembled genomes were performed using the ANIm (34) analysis on pyANI v0.2.12 (35). The genomes were dereplicated with drep v3.4.2 (18) using Mash (36) and MUMmer 3.0 (37), with a primary clustering ANI cutoff of 95% to distinguish between different species and a secondary clustering ANI cutoff of 99.5% to capture strain diversity to derive a representative set of genomes. The core genomes were determined using Roary (38) with an 80% BLASTp cutoff. PhyML 3.3 (39) trees of the core genome alignments of the representative genomes were constructed using the HKY85 substitution model with approximate likelihood ratio tests (aLRT). Trees were visualised on Geneious Prime 2024.0.7 (Biomatters). 16S rRNA alignments were performed using the Clustal Omega algorithm and visualised on Geneious Prime 2024.0.7 (Biomatters).

Pangenome analysis was performed using the anvi-pan-genome package and visualised on anvi’o v8 (20). Relevant gene clusters were extracted and functionally analysed with eggNOG-mapper v2 (21,22) to determine the respective COG categories. Metabolic pathways were identified using blastKOALA and KEGG Reconstruct (40,41). Phage defence systems were predicted using PADLOC v4.3 (30).

## Ethics statement

This study was conducted in accordance with the ethical principles of Good Clinical Practice and the Declaration of Helsinki. The local ethics committees of Unilever R&D (Port Sunlight, UK) approved the protocols before commencement of the studies and all subjects gave written informed consent.

## Data availability

Genomes of the 7 dereplicated genomes were deposited to the GenBank repository within the BioProject PRJNA1191350.

## Author contributions

R. H., R. C, B. M. and G. H. T. conceptualization; R. H., D. T. W, M. R., and B. M. validation; R. H., D. T. W., and G. H. T. formal analysis; R. H., D. T. W., I. K., J. H. and M. R. investigation; R. H. data curation; H. W., M. R. and M. H. resources; R. H. writing–original draft; R. H., D. T. W., S. M., I. K., H. W., A. J. W., B. M., and G. H. T. writing–review & editing; R. H. and S. M. visualization; A. J. W., B. M., and G. H. T. supervision; R. H., B. M., and G. H. T. project administration; B. M. and G. H. T. funding acquisition.

