## Supplementary Data for "High-resolution *Staphylococcus* profiling reveals intra-species diversity in a single skin niche"

```
Consensus                                 TATGGAGAGTTTGATCCTGGCTCAGGATGAACGCTGGCGGCGTGCCTAATACATGCAAGTCGAGCGAACAGACGAGGAGCTTGCTCCTCTGACGTTAGCGGCGGACGGGTGAGTAACACG   120
A1_1_H3st - 16S rRNA                      ........................................................................................................................   120
Staphylococcus_capitis_DSM_6717 - 16S rRNA ........................................................................................................................   120

Consensus                                 TGGATAACCTACCTATAAGACTGGGATAACTTCGGGAAACCGGAGCTAATACCGGATAACATGTTGAACCGCATGGTTCAACAGTGAAAGACGGTCTTGCTGTCACTTATAGATGGATCC   240
A1_1_H3st - 16S rRNA                      ........................................................................................................................   240
Staphylococcus_capitis_DSM_6717 - 16S rRNA ........................................................................................................................   240

Consensus                                 GCGCCGCATTAGCTAGTTGGTAAGGTAACGGCTTACCAAGGCAACGATGCGTAGCCGACCTGAGAGGGTGATCGGCCACACTGGAACTGAGACACGGTCCAGACTCCTACGGGAGGCAGC   360
A1_1_H3st - 16S rRNA                      ........................................................................................................................   360
Staphylococcus_capitis_DSM_6717 - 16S rRNA ........................................................................................................................   360

Consensus                                 AGTAGGGAATCTTCCGCAATGGGCGAAAGCCTGACGGAGCAACGCCGCGTGAGTGAAGAAGGTCTTCGGATCGTAAAACTCTGTTATTAGGGAAGAACAAATGTGTAAGTAACTATGCAC   480
A1_1_H3st - 16S rRNA                      ........................................................................................................................   480
Staphylococcus_capitis_DSM_6717 - 16S rRNA ........................................................................................................................   480

Consensus                                 GTCTTGACGGTACCTAATCAGAAAGCCACGGCTAACTACGTGCCAGCAGCCGCGGTAATACGTAGGTGGCAAGCGTTATCCGGAATTATTGGGCGTAAAGCGCGCGTAGGCGGTTTTTTA   600
A1_1_H3st - 16S rRNA                      ........................................................................................................................   600
Staphylococcus_capitis_DSM_6717 - 16S rRNA ........................................................................................................................   600

Consensus                                 AGTCTGATGTGAAAGCCCACGGCTCAACCGTGGAGGGTCATTGGAAACTGGAAAACTTGAGTGCAGAAGAGGAAAGTGGAATTCCATGTGTAGCGGTGAAATGCGCAGAGATATGGAGGA   720
A1_1_H3st - 16S rRNA                      ........................................................................................................................   720
Staphylococcus_capitis_DSM_6717 - 16S rRNA ........................................................................................................................   720

Consensus                                 ACACCAGTGGCGAAGGCGACTTTCTGGTCTGTAACTGACGCTGATGTGCGAAAGCGTGGGGATCAAACAGGATTAGATACCCTGGTAGTCCACGCCGTAAACGATGAGTGCTAAGTGTTA   840
A1_1_H3st - 16S rRNA                      ........................................................................................................................   840
Staphylococcus_capitis_DSM_6717 - 16S rRNA ........................................................................................................................   840

Consensus                                 GGGGGTTTCCGCCCCTTAGTGCTGCAGCTAACGCATTAAGCACTCCGCCTGGGGAGTACGACCGCAAGGTTGAAACTCAAAGGAATTGACGGGGACCCGCACAAGCGGTGGAGCATGTGG   960
A1_1_H3st - 16S rRNA                      ........................................................................................................................   960
Staphylococcus_capitis_DSM_6717 - 16S rRNA ........................................................................................................................   960

Consensus                                 TTTAATTCGAAGCAACGCGAAGAACCTTACCAAATCTTGACATCCTCTGACCCCTCTAGAGATAGAGTTTTCCCCTTCGGGGGACAGAGTGACAGGTGGTGCATGGTTGTCGTCAGCTCG  1080
A1_1_H3st - 16S rRNA                      ........................................................................................................................  1080
Staphylococcus_capitis_DSM_6717 - 16S rRNA ........................................................................................................................  1080

Consensus                                 TGTCGTGAGATGTTGGGTTAAGTCCCGCAACGAGCGCAACCCTTAAGCTTAGTTGCCATCATTAAGTTGGGCACTCTAAGTTGACTGCCGGTGACAAACCGGAGGAAGGTGGGGATGACG  1200
A1_1_H3st - 16S rRNA                      ........................................................................................................................  1200
Staphylococcus_capitis_DSM_6717 - 16S rRNA ........................................................................................................................  1200

Consensus                                 TCAAATCATCATGCCCCTTATGATTTGGGCTACACACGTGCTACAATGGACAATACAAAGGGTAGCGAAACCGCGAGGTCAAGCAAATCCCATAAAGTTGTTCTCAGTTCGGATTGTAGT  1320
A1_1_H3st - 16S rRNA                      ........................................................................................................................  1320
Staphylococcus_capitis_DSM_6717 - 16S rRNA ........................................................................................................................  1320

Consensus                                 CTGCAACTCGACTACATGAAGCTGGAATCGCTAGTAATCGTAGATCAGCATGCTACGGTGAATACGTTCCCGGGTCTTGTACACACCGCCCGTCACACCACGAGAGTTTGTAACACCCGA  1440
A1_1_H3st - 16S rRNA                      ........................................................................................................................  1440
Staphylococcus_capitis_DSM_6717 - 16S rRNA ........................................................................................................................  1440

Consensus                                 AGCCGGTGGAGTAACCTTTTGGAGCTAGCCGTCGAAGGTGGGACAAATGATTGGGGTGAAGTCGTAACAAGGTAGCCGTATCGGAAGGTGCGGCTGGATCACCTCCTT              1548
A1_1_H3st - 16S rRNA                      ............................................................................................................              1548
Staphylococcus_capitis_DSM_6717 - 16S rRNA ............................................................................................................              1548
```

**Supplementary Fig 1.** Full 16S rRNA alignment of YSMAA1_1_H3st and *S. capitis* DSM 6717 (type strain).

### *S. epidermidis* (scheme: sepidermidis)

| Isolate | ST | ArcC | AroE | Gtr | MutS | PyrR | TpiA | YqiL |
|---|---|---|---|---|---|---|---|---|
| YSMAS1_1_E3 | 297 | 1 | 2 | 2 | 2 | 2 | 1 | 3 |
| YSMAS1_1_E10 | 621 | 7 | 1 | 1 | 6 | 2 | 1 | 1 |
| YSMAS1_1_C8 | 225 | 1 | 13 | 7 | 2 | 2 | 1 | 29 |
| YSMAS1_1_C10 | 179 | 1 | 2 | 2 | 2 | 1 | 1 | 1 |

**Supplementary Table 1.** Multilocus sequence typing (MLST) allelic profiles of *S. epidermidis* isolates. Listed for each isolate are the detected sequence type (ST), allele types of each locus used in the relevant MLST schemes (www.pubmlst.org).

### *S. hominis* (scheme: shominis)

| Isolate | ST | ArcC | GlpK | Gtr | Pta | TpiA | Tuf |
|---|---|---|---|---|---|---|---|
| YSMAS1_1_A7 | 66 | 6 | 5 | 3 | 6 | 6 | 3 |
| YSMAS1_1_D4 | - | 18 | ~5 | 7 | 1 | 6 | 3 |

**Supplementary Table 2.** Multilocus sequence typing (MLST) allelic profiles of *S. hominis* isolates. Listed for each isolate are the detected sequence type (ST), allele types of each locus used in the relevant MLST schemes (www.pubmlst.org).

| Isolate | 1 | 2 | 3 | 4 |
|---|---|---|---|---|
| *S. capitis* YSMAA1_1_H3st | vga(A)LC, DQ823382, [lincomycin, clindamycin, dalfopristin, pristinamycin iia, virginiamycin m, tiamulin] | | | |
| *S. hominis* YSMAS1_1_A7 | mecA, BX571856, [amoxicillin, ampicillin, cefepime, cefixime, cefotaxime, cefoxitin, ceftazidime, ertapenem, imipenem, meropenem, piperacillin] | blaZ, NZ_JVAT01000021, [amoxicillin, ampicillin, piperacillin, penicillin] | fusC, KF527883, [fusidic acid] | |
| *S. hominis* YSMAS1_1_D4 | blaZ, CP003979, [amoxicillin, ampicillin, piperacillin, penicillin] | | | |
| S. epidermidis YSMAS1_1_C8 | fosB, CP000029, [fosfomycin] | | | |
| *S. epidermidis* YSMAS1_1_C10 | fosB, CP000029, [fosfomycin] | fusB, AY373761, [fusidic acid] | | |
| *S. epidermidis* YSMAS1_1_E3 | fosB, ACHE01000077, [fosfomycin] | fusC, KF527883, [fusidic acid] | msr(A), X52085, [erythromycin, azithromycin, telithromycin, quinupristin, pristinamycin ia, virginiamycin s] | mph(C), AF167161, [erythromycin, spiramycin, telithromycin] |
| *S. epidermidis* YSMAS1_1_E10 | fosB, CP000029, [fosfomycin] | fusB, AY373761, [fusidic acid] | | |

**Supplementary Table 3.** Full 16S rRNA alignment of YSMAA1_1_H3st and *S. capitis* DSM 6717 (type strain)
